# Harmonizing and integrating the NCI Genomic Data Commons through accessible, interactive, and cloud-enabled workflows

**DOI:** 10.1101/2022.08.11.503660

**Authors:** Ling-Hong Hung, Bryce Fukuda, Robert Schmitz, Varik Hoang, Wes Lloyd, Ka Yee Yeung

## Abstract

Cancer data is widely available in repositories such as the National Cancer Institute (NCI) Genomic Data Commons (GDC). These datasets could serve as controls or comparisons in compendium analyses with user data, avoiding the expense and time of generating additional datasets. However, the user must be able to process their new data in the same manner for these comparisons to be useful. This can be non-trivial. Although the executables themselves are usually available in repositories, the GDC pipelines that describe that entire analysis workflow are currently published as text-based standard operating procedures (SOPs). It is difficult to document a computational workflow to the level of detail and accuracy required to reproduce the results. Discrepancies between versions and exclusions of details accumulate as the documentation inevitably lags behind code revisions. We address this problem by converting the SOPs into a downloadable and executable format.

Specifically, we converted the GDC DNA sequencing (DNA-Seq) and the GDC mRNA sequencing (mRNA-Seq) SOPs into reproducible, self-installing, containerized, and interactive graphical workflows. These can be applied to reproducibly process user data and to harmonize datasets across repositories. Using our publicly available graphical workflows, we harmonize raw RNA-Seq datasets from the GDC and the Genotype-Tissue Expression (GTEx) project that were originally processed using different methodologies to illustrate the importance of uniform processing of control and treatment data for accurate inference of differentially expressed genes. By disseminating the analytical methodology in a reproducible and easily executed form, we greatly increase the utility of the GDC by enabling researchers to uniformly process custom data and datasets across multiple repositories to enhance data interpretation. Our approach and open-source executable workflows of making the analytical process as readily available as the data can be applied to other data repositories to increase their impact on scientific research.

## INTRODUCTION

Massive amounts of data are now available to enhance understanding and inform treatment of cancer. Large-scale programs such as The Cancer Genome Atlas (TCGA), Therapeutically Applicable Research to Generate Effective Treatment (TARGET) [1], and Clinical Proteomic Tumor Atlas Consortium (CPTAC) [2] have generated multi-omics data resources for diverse types of cancer. The National Cancer Institute (NCI) launched a new Cancer Research Data Commons (CRDC) website [3] to connect these diverse datasets with analytical tools in 2020. The CRDC provides access to different data-specific repositories, including the Genomic Data Commons (GDC) that stores raw sequencing data and derived results [4–6]. As of the v39.0 data release on December 4, 2023, the GDC consists of data from over 44 thousand cases across 79 projects [7]. Experimental strategies in the GDC include RNA sequencing (RNA-Seq), microRNA sequencing (miRNA-Seq), whole genome sequencing (WGS), whole exome sequencing (WXS), and targeted sequencing.

The CRDC provides different levels of data that reflect the amount of processing. Raw unprocessed sequence data (level 1) derived directly from patients may contain identifying information and are typically controlled access [8]. For the GDC, controlled access data requires dbGaP (database of Genotypes and Phenotypes) authorization and eRA Commons authentication. For data access, the GDC provides several methodologies. The simplest is the web-portal that allows users to browse and query the database using a graphical interface. However, the web-portal is not designed for downloading large datasets, such as raw sequence files which are typically in the range of 10-20 GB for GDC data [8]. For data download, the GDC has the standalone Data Transfer Tool which has an optional user interface (UI). However, this is an older tool that does not support the Gen3 authentication protocol which is the new standard for the CRDC databases. There is a newer Gen3 client that lacks the UI and is not documented on the GDC site. Finally, there is also a web API (Application Programming Interface) for programmatic access that can be used to download large datasets but requires technical expertise to use.

In contrast to raw sequence data, processed data such as transcript counts do not reveal the identity of patients and are often available with fewer restrictions. The datasets are smaller in size, and easier to use. However, methodologies for processing sequence data are heterogeneous and involve a wide variety of different software tools, parameters, and supporting data that are constantly being updated. For example, the GDC has published detailed documentation of analytic pipelines developed to process raw data [9, 10] including its workflow for analyzing RNA-Seq data using the STAR aligner [11]. While STAR is a popular aligner, HISAT [12], Bowtie [13], Kallisto [14], and Salmon [15] are examples of other aligners and pseudo-aligners that are frequently used for RNA-seq analyses. The GDC uses the Genome Reference Consortium Human Build 38 (GRCh38) with additional viral decoy sequences to increase the accuracy of alignments. While GRCh38 is in widespread use, differences in the masking and decoys give rise to slightly different reference sequences. The GDC uses GENCODE [16] annotations to map alignment coordinates to transcripts. GENCODE is not universally used with RefSeq [17] being a manually curated annotation alternative. Due to constantly improving technology and data, analytical pipelines can change between GDC releases. Early GDC data releases used the STAR aligner [11] that required two passes. Later releases used an improved version of STAR that aligned in one pass. Later data releases also use STAR to generate the transcript counts whereas earlier versions used HTSeq [18] to quantify counts. Earlier data releases use GENCODE v22 for annotations whereas the current release uses GENCODE v36.

Processing data using different pipelines affects results. Arora et al. compared well-established RNA-Seq processing pipelines using 6690 human tumor and normal samples from the TCGA and GTEx projects and reported major discrepancies in the abundance estimates that include disease-associated genes [19]. Arora et al. called for a community wide effort to develop gold standards to estimate mRNA abundances that could be used to harmonize data from different projects [19]. However, results that we present in this paper show that even minor changes in the software, versions, parameters, and supporting datasets can affect the identification of differentially expressed genes. Furthermore, these artifacts mitigate the effectiveness of a data standards approach for harmonization. Changes in methodologies and parameters are not capricious but reflect ongoing technological improvements. For example, we fully expect that new references and annotations derived from the new telomere to telomere human sequence will eventually be incorporated into GDC pipelines. Clearly, re-processing and harmonizing the entire repository upon every change in protocol is not a solution that scales with the rapidly growing amount of data. The dynamic solution that we propose is to facilitate and distribute the workflows so that they can be reproducibly applied to raw datasets and customized as methods, versions, and supporting data are updated. To accommodate the size of the raw datasets and the computational demands of the processing, the tool must be cloud enabled to minimize data transfers, and to take advantage of the enhanced throughput and scalable computational abilities afforded by the cloud. Additionally, the tool should be accessible and support interactive graphical analyses. Finally, the workflows should be portable, reproducible, and easily shared to allow researchers to reprocess custom user data or datasets from different repositories with identical software, versions, parameters, and supporting datasets. This will expand the usage and utility of large-scale data resources such as the CRDC.

### Our Contributions

In this manuscript, we present genomics workflows validated using data from the NCI Genomic Data Commons. These graphical, interactive, and cloud-enabled workflows are ready to be adopted to integrate data generated across different laboratories. Specifically, we have converted the text-based descriptions of the GDC data processing pipelines available at https://docs.gdc.cancer.gov/Data/Introduction/ to graphical workflows that are readily deployed from a public GitHub repository at https://github.com/BioDepot/GDC_Genomic_Workflows. In particular, we added GDC DNA sequencing (DNA-Seq)[10], the GDC mRNA-Seq workflows [9], as well as the Data Commons Framework Services (DCFS) Gen3 authentication [20] to provide integrated access to protected data from across the CRDC. Most importantly, these graphical workflows are dynamic. In other words, users can use a form-based user interface to customize these workflows by changing input parameters, updating versions of software, and providing annotations. We also demonstrate the utility of our workflows for harmonizing RNA-Seq data from TCGA and the Genotype-Tissue Expression (GTEx) [21] projects. Specifically, we demonstrate the impact of uniform re-processing of data versus direct use of processed RNA-Seq data on the inference of differentially expressed genes and present best practices for analyzing such data. Instead of developing static gold standard data processing pipelines for genomics data, we illustrate how our graphical workflows can be used to reproducibly distribute computational protocols that will enhance the flexibility, and ease of integration across multiple data sources.

### Related Work

In addition to enabling data sharing, the GDC provides software tools from the web-based data portal that supports data analysis, visualization, and exploration (DAVE) [22]. The NCI also supports the development of cloud-based platforms to analyze data hosted by the CRDC, including the Broad Institute’s Terra (formerly known as Firecloud) [23], the Institute for Systems Biology’s Cancer Gateway in the Cloud (ISB-CGC) [24, 25], and Seven Bridges Cancer Genomics Cloud [26, 27]. Both Terra and the ISB-CGC leverage Google Cloud to support cancer genomic analysis. Terra provides integrated access to the CRDC while providing pre-configured workspaces that support common use cases, such as the GATK best practices workflow. The ISB-CGC supports interactive web-based applications, Google Cloud APIs, and custom scripts and APIs for CRDC data access [24]. Seven Bridges is a commercial service using Amazon Web Services (AWS) or Google Cloud for bioinformatics analyses [26]. Users can drag multiple “apps” and parameters onto a canvas to connect them to define an executable workflow.

Most existing workflow execution platforms such as NextFlow [28] were designed around traditional batch workflows and scripting methods where a user interface such as DolphinNext [29] was appended afterwards. Other execution engines, such as Seven Bridges with integrated access to the CRDC, leverage on power tools for the Common Workflow Language (CWL), such as Rabix that supports editing of CWL scripts and visualization of CWL workflows. Galaxy is a web-server that provides a common web interface for users to create and execute workflows in a consistent hardware and software environment on a server or cluster [30]. While most Galaxy workflows are not containerized, Galaxy can use Bio-Docklets [31] to execute Docker workflows.

The Biodepot-workflow-builder (Bwb) [32] platform is an open-source, graphical platform for biomedical scientists to interactively execute workflows, monitor results, and adjust parameters. Each workflow defines an acyclic graph of executable modules (widgets) and the associated parameters. Upon loading a workflow, the Bwb application uses a browser or VNC client to display a set of connected graphical widgets, each of which represents a modular and containerized task. All commands defined in a Bwb workflow are executed inside a software container allowing for portable and reproducible execution on laptops, desktop servers, and across multiple cloud platforms. The containers are available through a public DockerHub and the Dockerfiles needed to create them are included with workflows. Unlike Galaxy that requires modifying a set of configuration files and scripts when importing tools and containers from non-Galaxy sources, Bwb provides specific GUI tools for customizing existing workflows and to facilitate the import of user scripts (in R, Python, CWL, WDL, Bash, Perl, and Java), and user defined Docker containers. Bwb workflows are saved as a directory of human readable text files that are made available and version controlled through a public GitHub. The open-source Bwb application is itself distributed as a public container. Features of Bwb include automated installation, form-based entry of parameters, and the ability to add new modules via drag-and-drop. Additionally, Bwb supports graphical output and interaction with applications that have a GUI such as Jupyter notebooks.

## RESULTS

### Graphical Genomics Workflows: Overview

We present a graphical and reproducible implementation of GDC genomics workflows in this section. In this work, all graphical workflows are implemented in the Bwb. Users can ***interactively*** start, stop, and modify these workflows through a drag-and-drop, point-and-click user interface in Bwb. Parameters in each module can be changed via a form-based user interface in Bwb. Bwb supports modules with their own graphical output and interfaces, including gnumeric spreadsheets[33], the Integrated Genome Viewer (IGV) [34], and Jupyter notebooks [35]. Since Bwb can export workflows as bash scripts of Docker commands, our GDC workflows can be run outside Bwb as bash scripts of containers or imported as containers in other workflow execution engines. We tested our workflows with controlled-access data from The Cancer Genome Atlas (TCGA) [36] and open-access data from the Cancer Cell Line Encyclopedia (CCLE) [37, 38] projects from the GDC. Our widgets, workflows, and documentation are available from our GitHub repository (https://github.com/BioDepot/GDC_Genomic_Workflows).

#### Integration with the Cancer Research Data Commons

The NCI Data Commons Framework Services (DCFS), powered by Gen3, is a set of software services that facilitate the hosting, management, and sharing of cancer datasets in the cloud[20, 39]. The “Fence” and “Arborist” services manage authentication and authorization so that controlled access data can be shared in the CRDC cloud infrastructure. There is a Gen3-client that interacts with these services and provides a command-line interface to upload and download files to and from a Gen3 data commons [20]. However, the client is not able to download all data-protected files. In this work, we created a widget that uses the Gen3-client to download files from the CRDC and uses the existing GDC web-api in the cases where Gen3 client fails. The download widget is portable due to containerization, uses graphical forms instead of a cryptic CLI, and can access all the files in the GDC.

To obtain access to controlled data in the GDC, researchers must first apply for access to specific projects or datasets through the NIH dbGaP, and then grant access to individuals in their labs. Some datasets, such as the Panel of Normals (PON) used in the DNA-Seq pipeline, that require controlled access but belong to no specific project. Currently, the Gen3 client can only download files that belong to a specific project and cannot download datasets such as PON. These datasets, however, can be downloaded using the GDC API. Consequently, we have consolidated these two methods into a Gen3 widget. **Figure 1** shows a screenshot of Bwb’s support for downloading controlled access files using the Gen3 client and the GDC API. The user authenticates Gen3 by signing into the dbGaP via the NIH eRA Commons to obtain the credentials file. Users can also provide an access token for the GDC API. This widget will attempt to use Gen3 to download the file (or manifest of files) and if that fails the widget will attempt to use the GDC API. This ensures that all controlled access files can be downloaded. Parallel downloads are supported using Bwb’s internal list-based scheduler if the user enters multiple files or manifests. Using this new widget, downloads of any controlled access dataset can now be incorporated into Bwb workflows, provided the user has the necessary dbGAP authorization. In particular, we used the widget in the GDC DNA-seq pipeline, to download the PON and sequencing read data from the GDC. A demonstration video for the Gen3 widget in the Bwb is available at https://youtu.be/8upzPouRGys.

**Figure 1.**
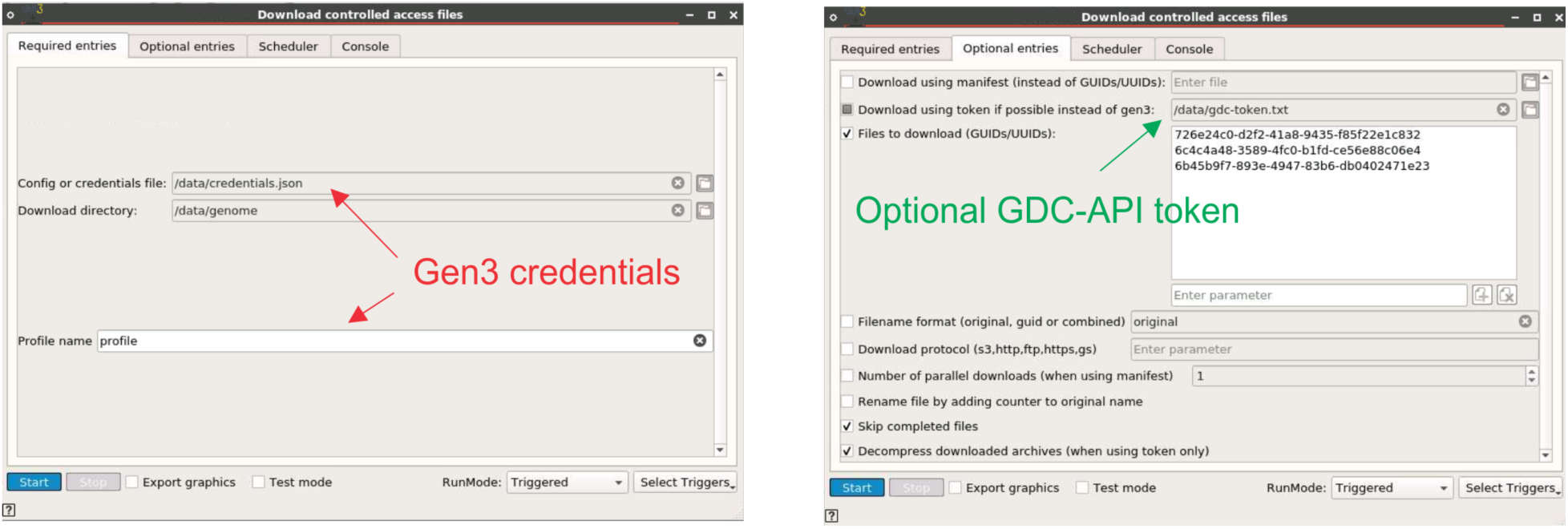
A screenshot of the panels from the Gen3 download widget. The required and optional entries panels from the Gen3 download widgets are shown. The user enters the location of the Gen3 credentials file and the desired profile to be used from the file. These are obtained by signing into the dbGaP database via the NIH eRA Commons. The user also has the option of entering the token file for use with the GDC API for files that cannot be currently downloaded by the Gen3 client. Multiple GUIDs or manifests can be entered, and parallel downloads are supported using Bwb’s built-in parallelism or using Gen3’s multithreaded downloading of manifests. We also include the option of decompressing the files on the fly as they are being downloaded.

#### mRNA-Seq workflow from the Genomic Data Commons

The GDC mRNA-Seq workflow [9] aligns raw sequence files to the GRCh38.d1.vd1 reference sequence using the STAR (Spliced Transcripts Alignment to a Reference) aligner [11], followed by the quantification step that outputs raw read counts and normalized read counts. In GDC Data Release versions 15 to 31, STAR [11] version 2.6.0c was used to compute the index and alignment, counts were obtained using HTSeq [18] using GENCODE [16] v22 as the reference annotation. Starting in GDC Data Release version 32, STAR version 2.7.5c is used with an additional input parameter, reference annotation are based on GENCODE v36, and counts obtained directly from STAR. This manuscript primarily focuses on Data Release version 32 since the documentation of the GDC mRNA-seq workflow [9] refers to this version extensively. **Figure 2 (a) and (b)** show screenshots of our implementation for GDC Data Release versions 15 and 32 in the Bwb respectively, consisting of the following steps: download the reference and sample data; create a genome index using the reference sequence, align reads to the reference, quantify the number of reads mapped to each gene, and calculate normalized gene expression values. The published GDC mRNA-Seq workflow includes the generation of gene fusion data using the STAR-Fusion v1.6 [9]. However, the CTAT genome libs from STAR-Fusion Release 1.6 is no longer available, so the gene fusion step is not automated in our v32 implementation. A demonstration video of the GDC mRNA-seq workflow (Data Release version 15) is available at https://youtu.be/YzFa9Een7Tc. An extended version of the mRNA-Seq workflow is shown in **Figure 3**, which the workflow includes harmonized uniform processing of TCGA and GTEx samples and Jupyter notebook widgets to perform differential expression analysis.

**Figure 2.**
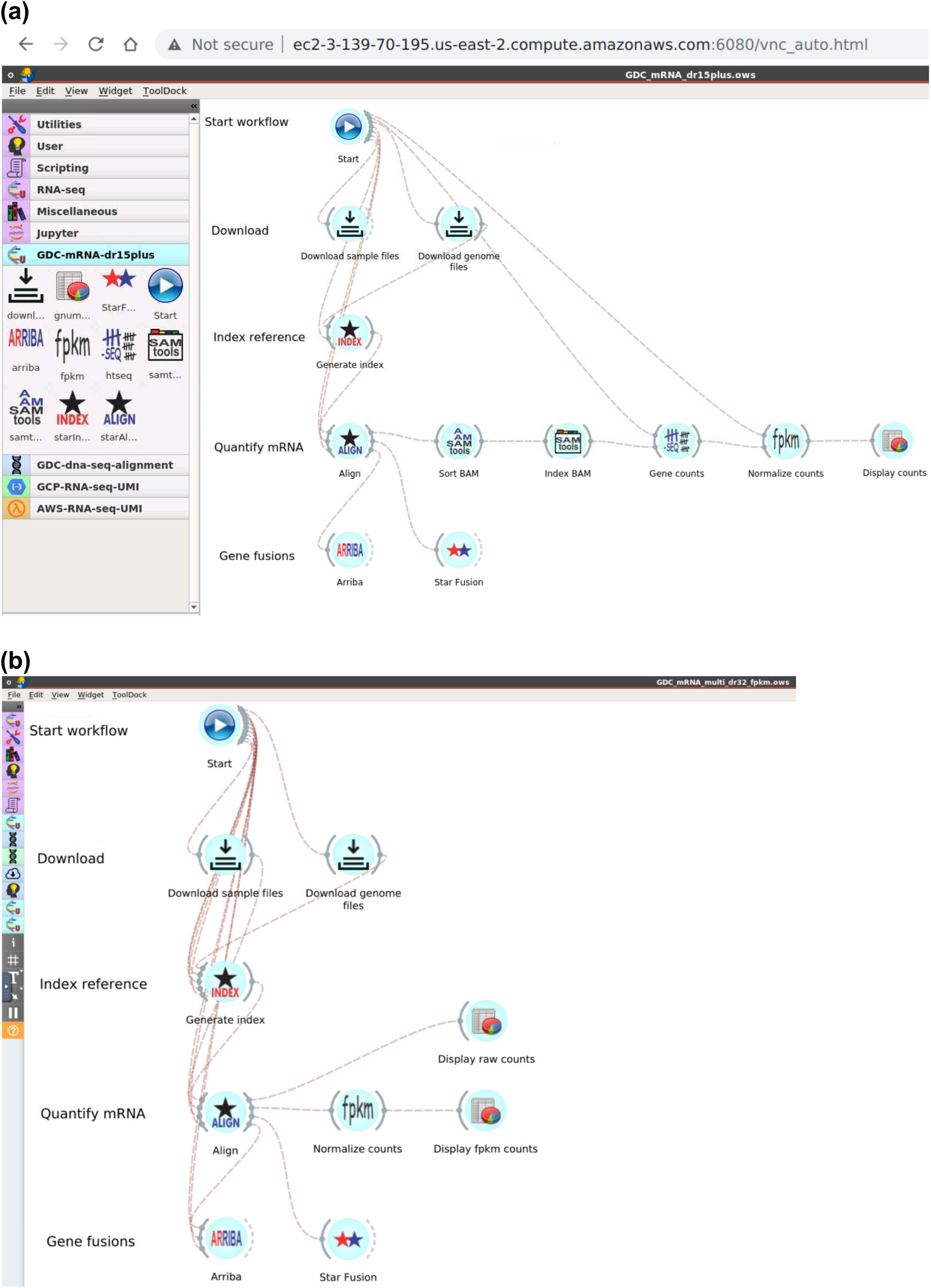
Screenshots of the GDC mRNA-Seq analysis workflows implemented in the Bwb. (a) GDC Data Release v15 mRNA-Seq workflow. (b) GDC Data Release v32 mRNA-Seq workflow. Each icon (widget) controls a separate containerized module. Double-clicking on a widget reveals graphical elements for parameter entry, starting and stopping execution, and displaying intermediate output. Lines connecting widgets indicate data flow between the execution modules. Connections and widgets can be added and removed using a drag-and-drop interface. The workflow itself is started by double-clicking on the start widget.

**Figure 3.**
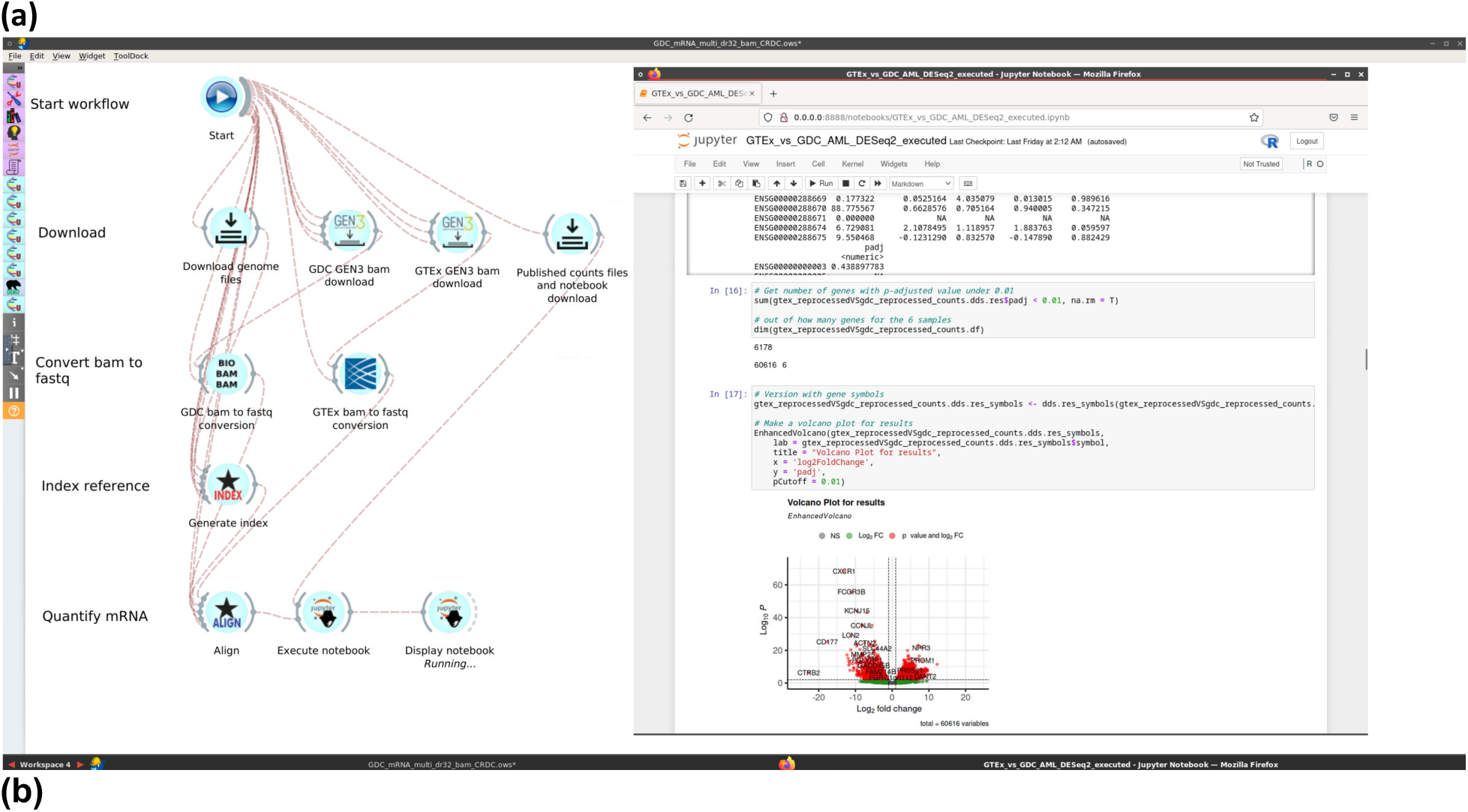
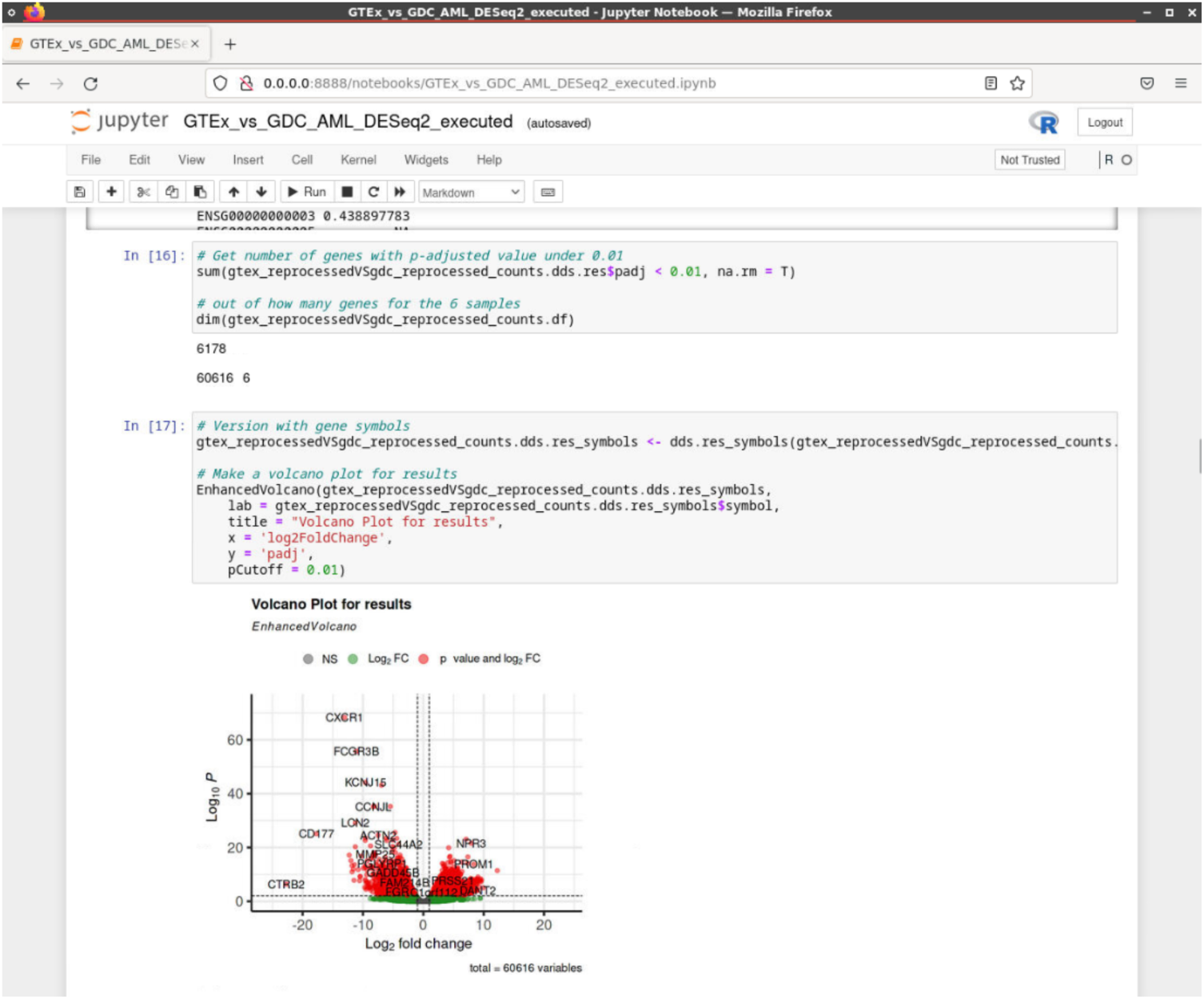
A screenshot of an extended mRNA-Seq analysis workflow implemented in the Bwb, and a screenshot of the Jupyter notebook after executing the workflow. (a) RNA-Seq workflow. Widgets are constructed with the same settings and parameters as the GDC Data Release v32 mRNA-Seq workflow. Gene fusion widgets are removed, GEN3 widgets to download BAM file samples for TCGA and GTEx are included, and widgets to convert downloaded BAM files into fastq file formats for harmonized uniform processing are added before running STAR. Outputs from STAR are analyzed for differential expression analysis at the end of the workflow using Jupyter notebooks. (b) Jupyter notebook with gene expression analysis is included in this integrated workflow. Count output files from STAR are used to perform differential expression analysis with DESeq2.

#### DNA-Seq workflow from the Genomic Data Commons

The GDC DNA-Seq workflows consist of six different methods for identifying somatic variants from WXS and WGS data from normal and tumor samples [10]. Our implementation in the Bwb, shown in **Figure 4**, consists of the following main steps: 1) download the reference and sample data; 2) convert BAM input files to fastq format; 3) align read groups to the reference genome using *bwa mem*; 4) perform variant calling using multiple callers; 5) annotate raw somatic mutations based on biological context and known variants from external mutation databases; 6) convert VCF to MAF files; and 7) display results in the Integrated Genome Viewer (IGV). A novel contribution from our team is that we have added functionality to create a batch file to load multiple variant files and regions of interest. A demonstration video of the GDC DNA-seq workflow is available at https://youtu.be/M7MCI83Q7_A.

**Figure 4.**
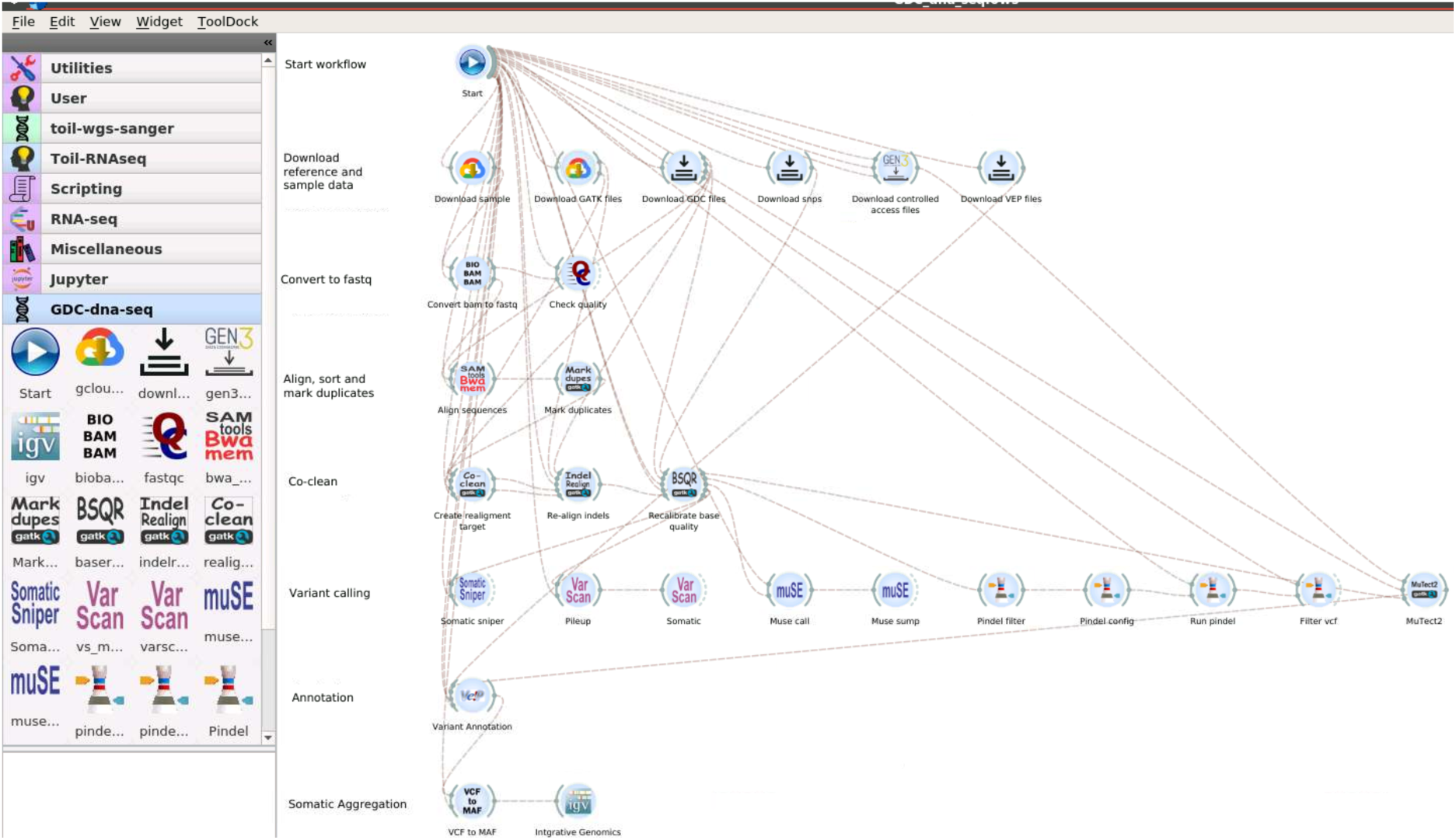
A screenshot of the GDC DNA-seq analysis workflow implemented in the Bwb. At the end of the workflow we have added an IGV widget which will automatically pop up with the regions of interest pre-loaded to allow the user to quickly evaluate the final MAF file.

### Harmonizing Cancer and Normal RNA-seq Data

In this section, we demonstrate the utility of our dynamic solution harmonizing RNA-Seq data from tumor samples in TCGA and normal tissue-specific samples in the Genotype-Tissue Expression (GTEx) [21] project. Specifically, we empirically studied the impact of different pipeline variations on the estimation of transcript abundance and inference of differentially expressed genes.

#### Data

For TCGA RNA-Seq data, we downloaded fastq files from the GDC Legacy Archive [40], BAM files, and counts data generated by the STAR aligner (Data Release version 32) from the GDC Data Portal at https://portal.gdc.cancer.gov/ [41]. Since the HTSeq files were removed from the GDC Data Portal starting in Data Release 32, we used patient case IDs to identify HTSeq count files’ case ID by cross-referencing to the manifest available from the GDC documentation GitHub [42]. This manuscript primarily focuses on Data Release version 32 to be consistent with the documentation of the GDC mRNA-seq workflow [9]. For the GTEx RNA-Seq data, we downloaded the v8 processed counts data from the GTEx Portal [43] and the controlled access BAM files from the AnVIL repository [44, 45].

#### Comparison of different GDC data releases

Since the RNA-Seq workflow had been changed substantially in Data Release version 32 compared to versions 15 to 31, our first step is to quantitatively compare the published raw counts from Data Release v15 HTSeq output counts file with the Data Release v32 STAR (v2.7.5c) output counts file. In particular, we downloaded published counts for sample TCGA-AB-2821 with case UUID: f6f9ed0d-2b3c-45b7-b214-853b5a207bac from the TCGA Acute Myeloid Leukemia (TCGA-LAML) project. After removing version numbers from the stable Ensembl gene IDs, there are a total of 56,485 common gene IDs in both count files. We observe that the published counts from GDC Data Release v15 are quite different from GDC Data Release v32, with 31,804 (56%) genes showing identical unnormalized counts. We computed the relative change ((v32 - v15)/v15) for each gene for which the v15 counts are non-zero. Among the 37,179 genes with non-zero counts, the median of the relative change is 0.0185, the 90^th^ percentile of the relative change is 0.5, and 55 genes show a relative change above 10. We then compared two other TCGA-LAML samples for their differences in counts between versions 15 and 32. Samples TCGA-AB-2828 (UUID: fc4ae4f8-f66b-4137-9821-e579b339cbf6) and TCGA-AB-2839 (UUID: cb262c7c-2646-45e3-bea9-376e48eefe65) both have the same total number of common genes between the two versions as TCGA-AB-2821 (56,485 genes). TCGA-AB-2828 has 31,729 (56%) genes with matching unnormalized counts between the two versions, and TCGA-AB-2839 has 31,079 (55%) genes with matching unnormalized counts. From calculating the relative change in TCGA-AB-2828, the median relative change among 35,583 genes with non-zero counts is 0.0279, the 90th percentile is 0.5882, and 57 genes exhibit a relative change greater than 10. For TCGA-AB-2839, the median relative change from 36,645 non-zero count genes is 0.02453, the 90th percentile is 0.6, and 79 genes exhibit a relative change greater than 10. **Table 1** shows the 8 genes with relative change above 100. This comparison illustrates that minor updates in data processing workflows could lead to major changes in the output counts of some genes. Our results highlight the need for a ***dynamic solution*** to re-process the raw data as workflows are being updated to adopt the latest version of aligners and annotation references.

**Table 1.** Genes with relative change ((v32 - v15)/v15) over 100 when comparing the GDC Data Release version 15 to version 32. Note that the stable Ensembl gene IDs are shown without the version numbers.

| Ensembl gene ID | gene symbol | GDC v15 counts | GDC v32 counts |
| --- | --- | --- | --- |
| ENSG00000157654 | PALM2AKAP2 | 7 | 2536 |
| ENSG00000197753 | LHFPL5 | 2 | 659 |
| ENSG00000245864 | MEF2C-AS2 | 1 | 147 |
| ENSG00000250891 | LINC02208 | 1 | 102 |
| ENSG00000253194 | AL137009.1 | 1 | 635 |
| ENSG00000260007 | AC107871.1 | 1 | 496 |
| ENSG00000265817 | FSBP | 1 | 157 |
| ENSG00000279170 | TSTD3 | 3 | 568 |

#### Comparison of published vs. reprocessed counts from GTEx

The GTEx project is a public resource profiling tissue-specific gene expression in non-diseased individuals [21]. The GTEx version 8 RNA-Seq processing workflow [46] used STAR v2.5.3a to align reads to the human reference genome GRCh38/hg38, based on the GENCODE v26 annotation. Subsequently, read counts and normalized TPM values were produced with RNA-SeQC v1.1.9 [47]. Thus the two projects used different versions of the aligner, different annotations, and probably most significantly, different software for obtaining read counts. Before we integrated the GTEx data with TCGA data, we studied the impact of the GTEx RNA-Seq pipeline versus the GDC RNA-Seq pipeline on unnormalized counts. Towards this end, we downloaded data from three whole blood samples, namely GTEX-N7MS-0007-SM-2D7W1 (abbreviated as “N7MS”), GTEX-NFK9-0006-SM-3GACS (abbreviated as “NFK9”), and GTEX-O5YT-0007-SM-32PK7 (abbreviated as “O5YT”). The publicly available v8 processed counts were downloaded from the GTEx data portal [43], while the BAM files were downloaded from AnVIL [44, 45]. We applied the GDC version 32 pipeline to process the downloaded BAM files from GTEx.

First, we computed the Pearson’s correlation coefficient between the published and reprocessed counts for each of these three whole blood samples. The correlation coefficients are 0.998, 0.993, and 0.995 respectively for each of the N7MS, NFK9, and O5YT samples. We observed that the number of non-zero published counts are 23,906, 22,086, and 23,997 respectively for each of these three samples out of a total of 55,617 genes. Next, we computed the relative change, defined as ((reprocessed - published)/published), for each gene with a non-zero published count. The medians for the relative change were 0.421, 0.420, and 0.439 respectively for each of these three blood samples. The 99^th^ percentile of the relative changes were 4, 3.25, and 4 respectively. As a control, we compared the three published samples to the three reprocessed samples by applying DESeq2 [48], and observed that 625 genes have adjusted p-values under 0.05. To summarize, while the correlation coefficients between the published and reprocessed counts are high, there are some genes with substantial changes when the raw data was reprocessed with the GDC workflow. This empirical study highlights the need to harmonize the RNA-Seq data using the identical workflow before integration.

#### Integration of cancer and normal RNA-seq data by reproducibly sharing dynamically updated workflows

We next illustrate the importance of harmonizing data from different repositories and demonstrate how this can be accomplished using Bwb workflows for a real-world application, Specifically, we used our GDC Data Release v32 RNA-Seq workflow to integrate tumor data from the GDC and normal data from the GTEx project. We downloaded BAM files from three cases (TCGA-AB-2821, TCGA-AB-2828, TCGA-AB-2839) in the TCGA-LAML project. We harmonized the transcript abundance of these tumor samples and the three whole blood normal samples discussed in the previous sub-section using our implementation of the GDC version 32 workflow. Subsequently, we used DESeq2 [48] to infer differentially expressed genes. Out of the 60,616 Ensembl gene IDs, 6178 gene IDs show an adjusted p-value under 0.01. We then mapped the Ensembl gene IDs to gene symbols using biomaRT [49]. **Figure 5** shows the volcano plot of the DESeq2 output. We repeated the analysis using data that was not harmonized. We concatenated the v8 published counts from GTEx and v32 published counts from GDC, applied DESeq2, and recorded the top 10 differentially expressed genes. **Table 2** compares the differentially expressed genes inferred from concatenation of published counts versus those inferred from harmonized uniform GDC re-processing. We observe that CXCR1 has the most significant (smallest) adjusted p-values in both scenarios with zero change in rankings. As another example, FCGR3B is the second most significant differentially expressed gene in the harmonized re-processed scenario, with the reduction of one rank in the concatenation of published counts scenario. We observe that 5 of the top 10 (CD68, GPS2, ARL6IP4, GABARAP, CHKB) differentially expressed genes in the concatenation of published counts scenario exhibit dramatic change in rankings (over 10,000). In other words, half of the top 10 differentially expressed genes could be bogus if we don’t reprocess the raw RNA-Seq data using uniform pipelines. This example illustrates the importance of harmonized uniform processing of RNA-Seq data.

**Figure 5.**
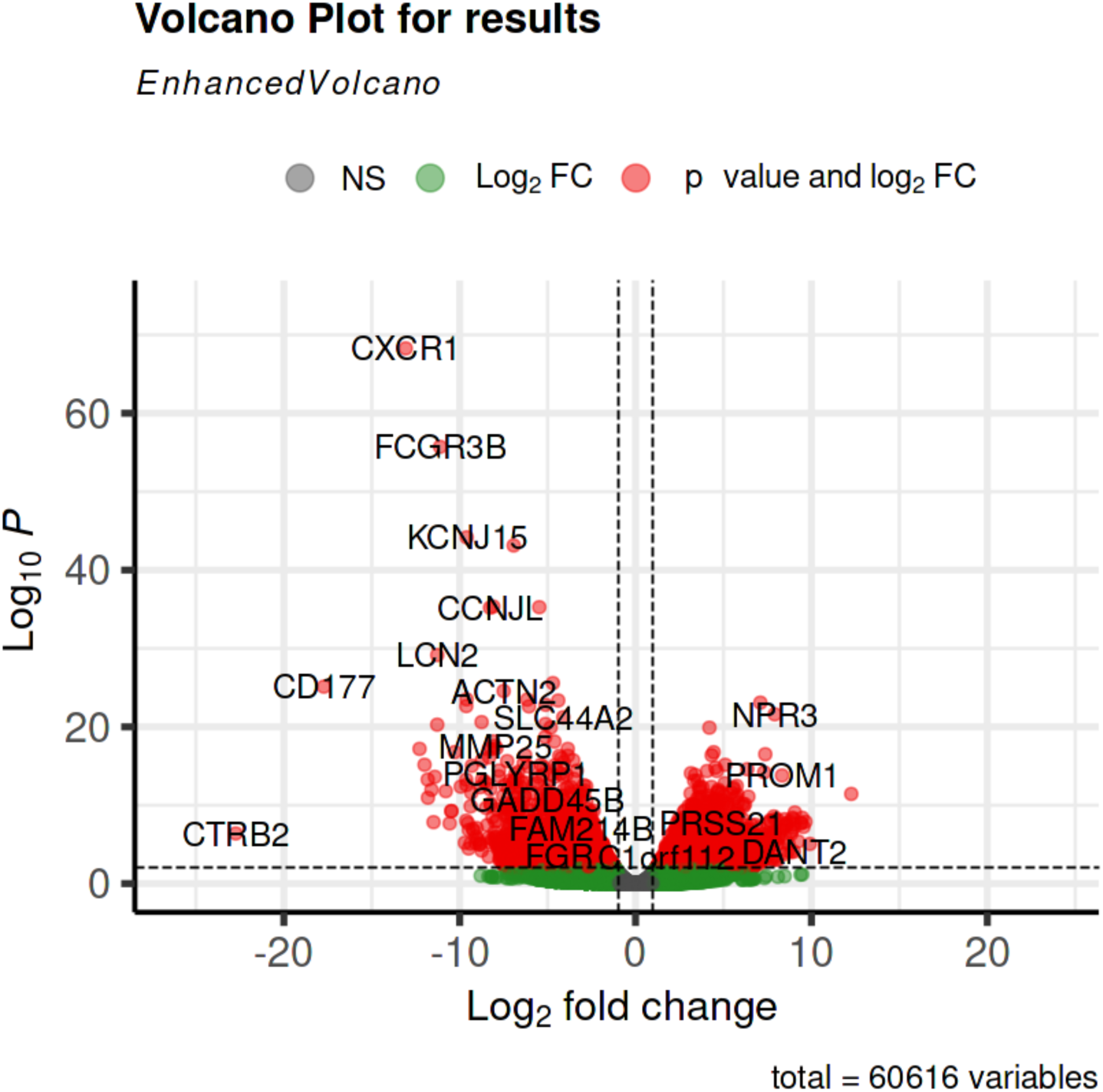
Volcano plot for differential genes comparing cancer vs. normal blood samples.

**Table 2:** Comparison of the top 10 differentially expressed genes inferred from concatenation of published counts (“published vs published”) versus those inferred from harmonized uniform GDC re-processing (“reprocessed vs reprocessed”). The column “rank delta” corresponds to the change in rank. In particular, a zero-rank delta means no change in rank, e.g. CXCR1. The column “padj” shows adjusted p-values obtained from DESeq2.

| <u>Published vs</u> |  |  | <u>Reprocessed vs</u> |  |  |
| --- | --- | --- | --- | --- | --- |
| <u>Published</u> |  |  | <u>Reprocessed</u> |  |  |
| padj | Gene | rank $\Delta$ | rank $\Delta$ | Gene | padj |
| 5.98E-70 | CXCR1 | 0 | 0 | CXCR1 | 5.26E-69 |
| 2.67E-57 | CD68 | 26180 | 1 | FCGR3B | 1.91E-56 |
| 1.66E-56 | FCGR3B | -1 | 4 | KCNJ15 | 6.68E-45 |
| 2.33E-56 | GPS2 | 20906 | 5 | FAM157A | 7.42E-44 |
| 7.13E-56 | ARL6IP4 | 19617 | 13 | TREML3P | 4.54E-36 |
| 5.24E-51 | RNASEK | 379 | 15 | R3HDM4 | 5.45E-36 |
| 6.39E-46 | KCNJ15 | -4 | 10 | CCNJL | 6.41E-36 |
| 3.34E-45 | GABARAP | 10206 | 16 | LCN2 | 7.05E-30 |
| 1.31E-43 | FAM157A | -5 | 35 | YPEL3 | 2.65E-26 |
| 9.77E-43 | CHKB | 26113 | 22 | CD177 | 7.51E-26 |

#### Best practices when processing RNA-seq data

Our proof-of-concept integration of TCGA and GTEx RNA-seq data illustrates the importance of uniform data processing starting from raw sequence data with the same workflow, same input parameters, and the same versions of software tools and annotations. The graphical, pre-configured and easily updatable workflows presented in this paper can be used to uniformly process raw sequence data generated by different laboratories or across different projects. In particular, GDC RNA-seq and DNA-seq workflows with integrated access to the NCI Genomic Data Commons are presented. In addition to running the GDC RNA-seq workflow starting from the raw fastq input files, we have also executed the RNA-seq workflow using BAM input files. There are three different BAM files listed for each case ID in the GDC Data Portal: chimeric, genomic, and transcriptomic. We experimented with the conversion from BAM files to fastq using different parameters in Biobambam [50], Samtools [51], and Picard [52], and observed that the genomic BAM files converted using Biobambam with exclude parameter set to off and Samtools produced the sequences in the fastq file provided by the GDC Legacy Archive [40]. This conversion from BAM to fastq is implemented and included in our GDC RNA-seq workflows.

## DISCUSSION AND CONCLUSIONS

Using RNA-Seq data as a case study, we demonstrate the need to reprocess raw sequencing data since published counts change over different data releases with updated versions of aligners and reference annotations. We also show the need to harmonize raw sequencing data generated by different projects by re-processing the data with the same RNA-Seq workflow. Our observations echo the findings by Arora et al. [19]. However, instead of calling for a concerted, community-wide gold-standard for data processing, we provide ***a dynamic solution*** that distributes the computational methodology in a reproducible, accessible, customizable, and cloud-enabled manner to facilitate the reprocessing of data.

Our open-source graphical workflows are containerized and ready to be deployed on any cloud platform or local host with Docker installed. These cancer genomic workflows implement the SOPs published by the NCI Genomic Data Commons (GDC). Our workflows leverage the NCI DCFS Gen3 framework to enable integration of controlled access data from the Cancer Research Data Commons. Due to the modular nature of the Bwb (i.e. each module is encapsulated in a software container), these GDC workflows can be customized and adapted using a graphical user interface. These workflows can also be exported as bash scripts and software containers and can be deployed outside the Bwb platform. New widgets can be rapidly added and updated using the form-based user interface without writing additional GUI code. We demonstrate the utility of our graphical workflows for harmonizing RNA-Seq data from TCGA and the Genotype-Tissue Expression (GTEx) [21] projects.

## MATERIALS AND METHODS

### Implementation of GDC genomic workflows

Starting from the text description of the workflows available on the GDC website, we identified the component scripts/executables, versions, and parameters. We then decomposed the workflows into individual self-contained data-processing modules. For each module, we built Docker containers, and uploaded them to DockerHub. Graphical widgets were constructed in Bwb and connected to form the complete workflows. In addition, we added a “Start” widget to specify the directory structures and other global parameters. The connections between the widgets indicate and control the dataflow, dependencies, and sequence of execution. Thus, each step in our workflows is encapsulated in a Docker container, with specific version tags to ensure software dependencies and compatibility. The modular approach facilitates re-use and customization of the workflows and their components.

## AVAILABILITY OF SUPPORTING DATA AND MATERIALS

- Project name: GDC Genomics Workflows
- Project homage page: https://github.com/BioDepot/GDC_Genomic_Workflows
- Contents available for download: All workflows reported in this manuscript. Demonstration videos, Jupyter notebooks, documentation, Dockerfiles, scripts.
- Operating System(s): Linux. Also tested on AWS.
- Programming language(s): Python, Bash
- License: MIT license

## DECLARATIONS

## Consent for publication

Not applicable.

## Competing Interests

LHH and KYY have equity interest in Biodepot LLC, which receives compensation from NCI SBIR contract numbers 75N91020C00009 and 75N91021C00022. The terms of this arrangement have been reviewed and approved by the University of Washington in accordance with its policies governing outside work and financial conflicts of interest in research.

## Author’s contributions

Conceptualization and funding acquisition: L.H.H. and K.Y.Y. Software and investigation: L.H.H., R.S., B.F. and V.H. Cloud resources: L.H.H., W.L., and R.S. Formal analysis: B.F. Methodology: K.Y.Y. and L.H.H. Visualization: L.H.H. and B.F. Validation: L.H.H. Supervision: L.H.H. W.L. and K.Y.Y. Writing - original draft preparation: K.Y.Y. Writing - reviewing and editing: All authors. Project administration: K.Y.Y. The authors read and approved the final manuscript.

## Acknowledgements

LHH, WL, and KYY were supported by NIH grant R01GM126019, NCI SBIR contracts 75N91020C00009 and 75N91021C00022. RS, BF and VH were supported by NCI SBIR contracts 75N91020C00009 and 75N91021C00022. LHH and KYY are also supported by NIH grants U24HG012674 and R03AI159286. The funding bodies played no role in the design of the study and collection, analysis, and interpretation of data and in writing the manuscript.

The content is solely the responsibility of the authors, and does not necessarily represent the official views of the National Institutes of Health.

We would like to acknowledge cloud credits from Amazon Web Services and IBM Cloud.

## List of Abbreviations

AMI: Amazon Machine Image
API: application programming interface
AWS: Amazon Web Services
Bwb: Biodepot-workflow-builder
CPTAC: Clinical Proteomic Tumor Atlas Consortium
CRDC: Cancer Research Data Commons
DCFS: Data Commons Framework Services
dbGaP: database of Genotypes and Phenotypes
DNA-Seq: DNA sequencing
DTT: Data Transfer Tool
EC2: Elastic Compute Cloud
GDC: Genomic Data Commons
IGV: Integrated Genome Viewer
miRNA-Seq: micro RNA sequencing
NCI: National Cancer Institute
NGS: Next-generation sequencing
PON: Panel of Normals
RNA-Seq: RNA sequencing
TARGET: Therapeutically Applicable Research to Generate Effective Treatment
TCGA: The Cancer Genome Atlas
WGS: whole genome sequencing
WXS: whole exome sequencing

## Notes

### Summary of Updates

In this revision, we updated the content to reflect the latest data releases from the NCI Genomic Data Commons. We also made our contributions in this work clearer by revising the title, abstract and introduction. In addition, we re-tested our workflows, cleaned up the GitHub repository, added documentation, and include only the workflows that work.

